# Chemical probe-based Nanopore Sequencing to Selectively Assess the RNA modifications

**DOI:** 10.1101/2020.05.19.105338

**Authors:** Soundhar Ramasamy, Vinodh J Sahayasheela, Zutao Yu, Takuya Hidaka, Li Cai, Hiroshi Sugiyama, Ganesh N. Pandian

## Abstract

Current methods to identify RNA modifications with short-read sequencing are laborious and direct RNA sequencing gets proclaimed as the viable alternative. Herein, we harness the selective reactivity of the acrylonitrile towards the Inosine (**I)** and pseudouridine (**Ψ**) modifications and developed a chemical probe-based direct RNA sequencing method. We first demonstrated that the chemical probe-induced differential signature profile using nanopore sequencing could facilitate the selective assessment of **I** and **Ψ** in the *in vitro* synthesized RNA. Furthermore, we verified the **I** and **Ψ** modification with single-nucleotide resolution using RNA derived from mouse brain without the need for a null dataset using knockouts. Our chemical probe-based nanopore sequencing strategy can be extended to profile multiple RNA modifications on a single RNA and may facilitate the diagnosis of disease-associated epitranscriptome markers by generating a comparative dataset in clinical scenarios.

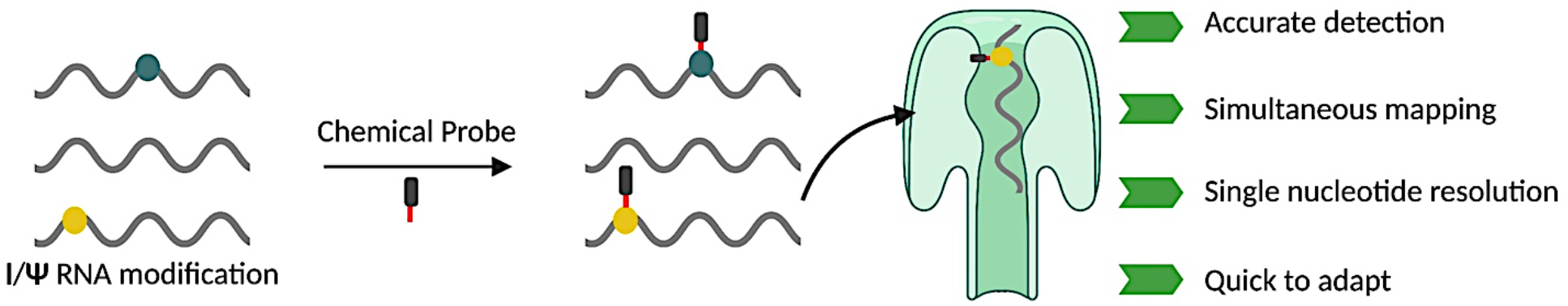

---

RNA modifications exert a spectrum of regulatory function over mRNA, ranging from base pairing^1^, structure^2,3^, stability^4,5^, decay^4,6^, translation^7,8^, microRNA binding^9,10^, and codon potential^8^. To date, 172 modifications (Modomics)^11^ collectively termed as “Epitranscriptome” are known to exist in biological systems. Transcriptome profiling methods have revealed the biological significance of the RNA modifications in complex diseases like cancer and neurodegenerative disorders. Most of the existing techniques use the short-read sequencing (SRS) platform, which has high accuracy and coverage. However, RNA modifications are silent to the reverse transcription used in SRS library preparation. Many protocols employ antibody and modification-specific chemicals for adduct generation^12^. These adduct induced mutation or truncation profiles are used as a proxy identifier of modifications with single-nucleotide resolution. The general limitations of the current methods are a) laborious in sample preparation, b) lesser reproducibility across replicates due to multi-step sample preparation, c) extracting stoichiometry of modifications are error-prone because of RNA fragmentation during SRS library preparation, and d) difficult to simultaneously map multiple modifications in single run^13^. Consequently, there is an increasing demand to develop a “Quick to adapt” RNA modification profiling methodologies that can be extended and applied in the clinical situation, where the input RNA is limited and RNA modification-associated enzyme knockouts are not possible.

Direct RNA-Seq (dRNA-Seq) using Oxford nanopore technologies obviates the need for reverse transcription and overcomes the above-described shortfalls with a unique sequencing strategy by ratcheting DNA/RNA into a protein pore. The migrating nucleic acid triggers a distinct change in the stable current maintained across the pore and the resulting pattern is basecalled to a corresponding nucleic acid sequence with machine learning algorithms. Because dRNA-Seq directly uses the native RNA, their current pattern and mapped reads reveal RNA modifications primarily through mismatch errors and by the alterations in intensity, dwell time or deletion parameter. Above ONT parameters are readily available from dRNA-seq reads even without the need of any customized treatment on native RNA^14,15^. However, the RNA modification associated mismatch errors can also be misinterpreted with that from single-nucleotide polymorphism/variations (SNP/SNV) or the intrinsic ONT platform noise. Accordingly, Liu et al., devised a strategy to identify N6-methyladenosine (**m**^**6**^**A)** RNA modification associated mismatch errors^15^ by comparing the wild type RNA with modifications and contrasting them with the modification-free RNA derived from the knockout studies. However, this strategy was demonstrated in the model organisms and cannot be applied to clinical scenarios in humans where knockouts are impossible. Also, RNA modification crosstalk will result in non-homeostatic epitranscriptome landscape in knockouts i.e., **m**^**6**^**A** knockouts are shown to have increased inosine (**I**) modification^16^, which may again present itself as a mismatch in dRNA-Seq. Thus, there can be an underestimation of **m**^**6**^**A** modification sites when scoring for the absence of mismatch errors between **m**^**6**^**A** wild type and its knockouts. Therefore, a strategy to differentiate **m**^**6**^**A** and **I** modification sites, particularly in mammalian systems, will be useful. Here we harness the selective reactivity of acrylonitrile towards different RNA modifications and demonstrate a proof-of-concept study to artificially induce differential mismatch errors that facilitate the selective assessment of **I** and pseudouridine (**Ψ)** mismatch errors from other noises.

Acrylonitrile selectively cyanoethylates **I** and **Ψ** at N1 position to form N1-cyanoethyl **I** (**CEI**) and N1-cyanoethyl **Ψ** (**CEΨ**), respectively **(Figure 1A)**^17^. Between the above-mentioned adducts, **CEI** stalls the reverse transcriptase during cDNA synthesis, resulting in truncated short reads. In comparison, **CEΨ** adduct remains silent and undetectable. Hence acrylonitrile-based **I** chemical erasing sequencing only detects **I** modification in SRS platform^18,19^. Herein, our notion is that **CEI**/**CEΨ** dRNAseq parameters will be different from that of **I/Ψ**, thereby facilitating RNA modification detection from dRNA-Seq reads, without the need for knockout derived RNA. Furthermore, as shown in our workflow (**Figure 1B**), acrylonitrile’s dual reactivity towards **I** and **Ψ** could also facilitate the selective assessment of both these biologically significant modifications. To comprehend the alterations in dRNA-Seq parameters, we first performed optimization studies by generating synthetic RNA using in vitro transcription (IVT) reaction with the following three conditions unmodified, modified [**I, Ψ** and **m**^**6**^**A**] and acrylonitrile treated (**CEI**/**CEΨ**) (**Figure 1B and S1A**). Because both **m**^**6**^**A** and **I** converge on the adenine(**A**) nucleobase, **m**^**6**^**A** negatively regulates **A**-to-**I** modification. A comparative assessment to differentiate **m**^**6**^**A** from **I** based on their unique dRNA-Seq profile was performed. We generated modified RNAs by replacing conventional NTPs with modified nucleotides in IVT. For **I** RNA modification, we synthesized two variants of **I** modified RNA (100% and 50%) transcripts. During dRNA-Seq, the 100% modified transcript did not yield quality reads upon acrylonitrile treatment, which suggests an unfavorable interaction between the pore and the transcript. On the other hand, such an issue in read quality was not observed in the 50% modified RNA transcript. Therefore, we used 100% and 50 % modified transcripts for **I** and **CEI** conditions, respectively. The **I** and **Ψ** modified RNA’s were subjected to cyanoethylation reaction with acrylonitrile treatment for 30 mins and the reactivity was confirmed through HPLC and mass spectrometry (**Figure S2 and S3)**. The size and concentration of all RNA’s were confirmed using a bioanalyzer (**Figure S4)**.

**Figure 1.**
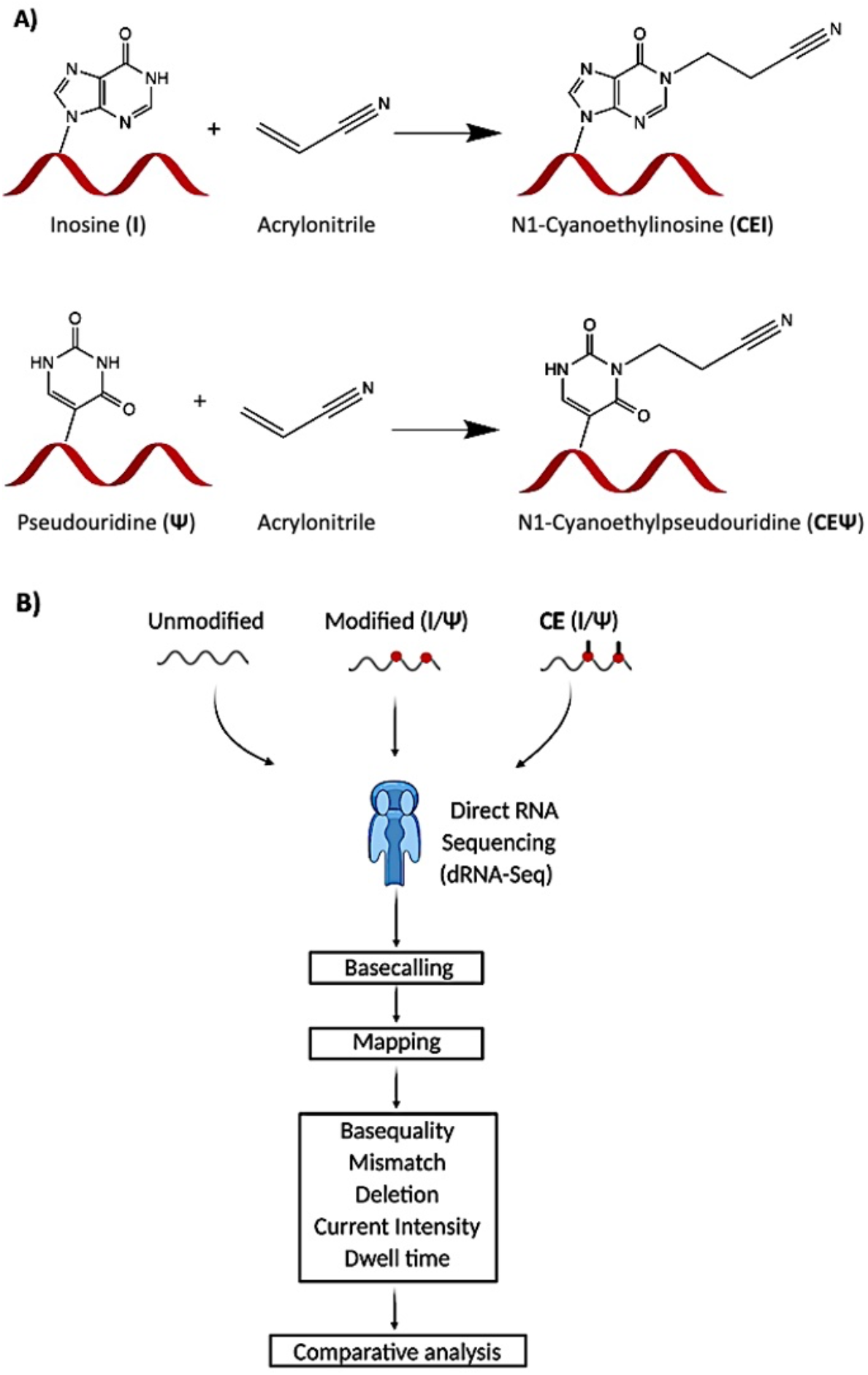
A) Schematics of acrylonitrile cyanoethylation reaction. Acrylonitrile selectively reacts with Inosine (**I**) and Pseudouridine (**Ψ**) in the RNA to form N1-cyanooethylinsoine (**CEI**) and N1-cyanoethylpseudouridine (**CEΨ**), **B)** Overview of the workflow used in this study to identify **I** and **Ψ** using selective chemical probe by Nanopore sequencing

We then performed dRNA-Seq using unmodified (UM), modified **(I, Ψ, m**^**6**^**A)**, and acrylonitrile treated (**CEI**/**CEΨ**) RNA transcripts and analyzed the chemical probe induced mismatch errors. While the UM transcripts showed almost no mismatch errors, the modified transcripts displayed consistent mismatch errors on the sites incorporated with modifications. (**Figure 2A and 3A)** Also, a remarkable difference in the error pattern was observed for all three modifications where mismatch errors for **I** was confused between **A**/**G** (**Figure 2B, 2C, and S5A) and Ψ** were confused mostly between **C**/**U** (**Figure 2B, 3B, and S5B)**. For **m**^**6**^**A**, the mismatch error profile was dominated with **A** (**Figure S6**) over other nucleotides. Other dRNA-Seq parameters like base quality, signal intensity, dwell time and deletion were also altered between UM and modified regions **(Figure 2E-H, Figure 3 E-H, and Figure S6 D-G)**. Taken together, all three modifications in IVT RNA could be selectively assessed in dRNA-Seq as mismatch errors with a unique signature profile. Subsequently, we compared the mismatch errors between modified and acrylonitrile-treated conditions to validate our hypothesis to identify RNA modifications using chemical probe-induced differential mismatch errors. The results indicated that the mismatch errors for **I** that was confused between **A**/**G** were shifted mostly towards **G** in **CEI** condition. **(Figure 2C and 2D**). On the other hand, the mismatch errors for **Ψ** that was confused between **C**/**U** displayed only a slight shift towards **C** (**Figure 3C and 3D**). This remarkable shift in the chemical probe induced mismatch error pattern generated by **CEI** can be attributed to acrylonitrile’s higher reactivity towards **I** than **Ψ (Ref 17)**. We then evaluated two ONT analysis packages (TOMBO and Nanocompore) for their ability to optimally assess RNA modifications by comparing modified (**I**/**Ψ**) and acrylonitrile treated (**CEI**/**Ψ**) reads. Both packages follow a similar workflow where the first step is resquiggle; the raw signal is matched to the reference sequence file, and then signals are compared between the control (modified) with the test (acrylonitrile treated) to identify the regions with a statistically significant signal difference (Δ signal). The results indicated that TOMBO could recover all modified sites in both **I** and **Ψ**, (**Figure 2I**,**3I, S7 and S8**). On the other hand, Nanocompore revealed sites only in **I** vs **CEI** and not in **Ψ** vs **CEΨ** (data not shown). Therefore, we employed TOMBO for all further analysis where the signal level at each reference position was compared between the control and test sample. The statistics of Δ signal is scored by default using the K-S test, whose statistics score ranges from zero to one, with values moving towards one indicating regions with a significant Δ signal. As shown in **Figure 2I**, Δ signal statistical significance was mostly scored by TOMBO over the modification containing regions. In a few regions, some statistically significant peak value falling maximum five nucleotides away from the actual modification site was observed, and this could be attributed to nanopore K-mer based signal handling. (**Figure 3I**).

**Figure 2.**
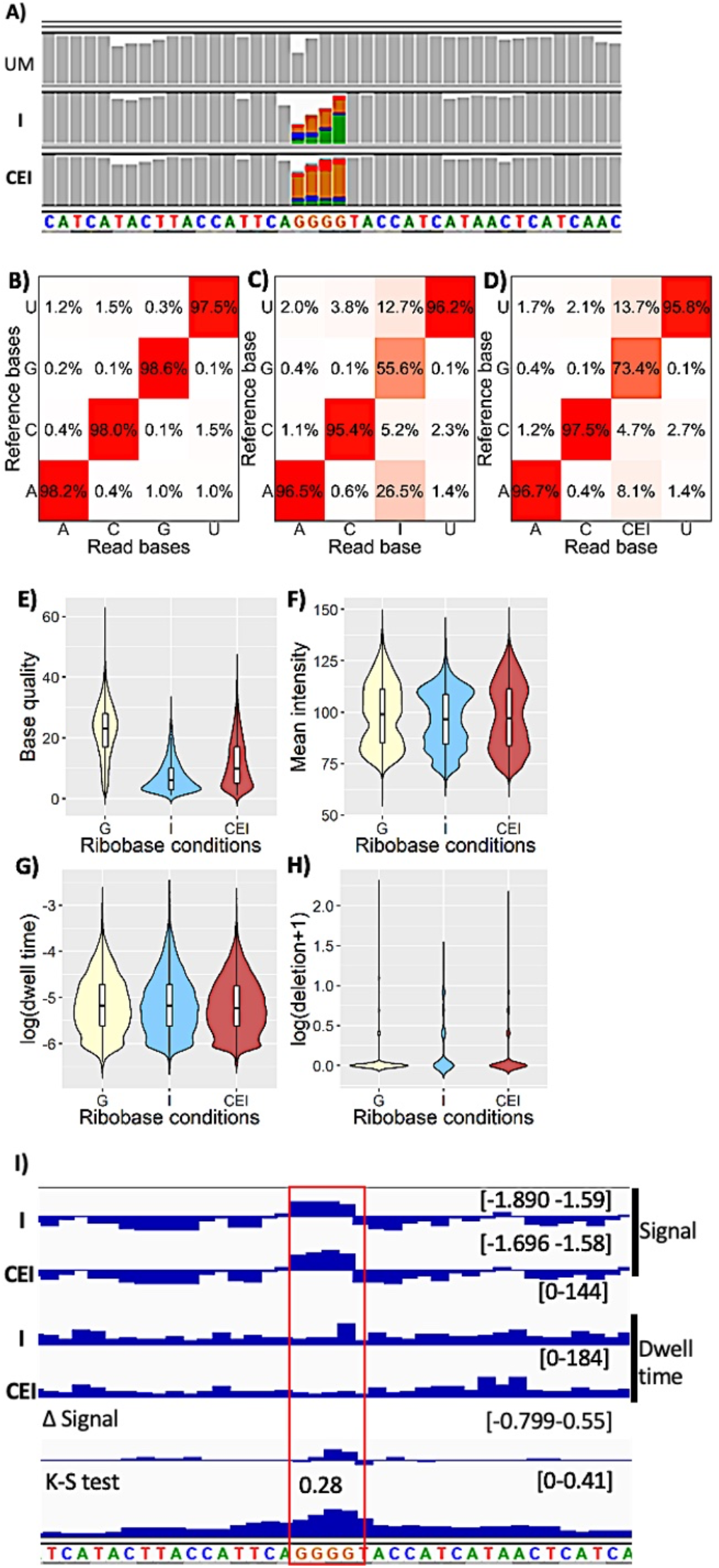
Altered nanopore parameters by I and CEI modified nucleobases. **A)** IGV snapshot of unmodified (UM), Inosine (I) and cyanoethyl inosine (CEI) transcripts showing mismatch. Mismatch frequency > 0.2% are represented in colors. Green(adenosine), orange (guanosine), blue (cytosine) and red (thymine). **B, C & D**) Substitution matrix of UM, **I** and **CEI** transcripts native reads, respectively. The x-axis represents the base identity of nanopore reads. The y-axis represents base identity of reference transcript **E-H**) Violin plot showing kernel density estimate & inner boxplot showing interquartile range and median of unmodified, **I** and **CEI** nucleobase. **E)** Base quality, **F)** Mean intensity, **G)** Dwell time and **H)** Deletion parameters. **I)** IGV snapshot showing signal comparison between **Ψ** (control) and **CEΨ** (samples) using TOMBO. The K-S test was used to compare signal difference ((Δ signal) between **Ψ** and **CEΨ** conditions. The K-S test value > 0.2 is considered to harbor statistically significant signal difference.

**Figure 3.**
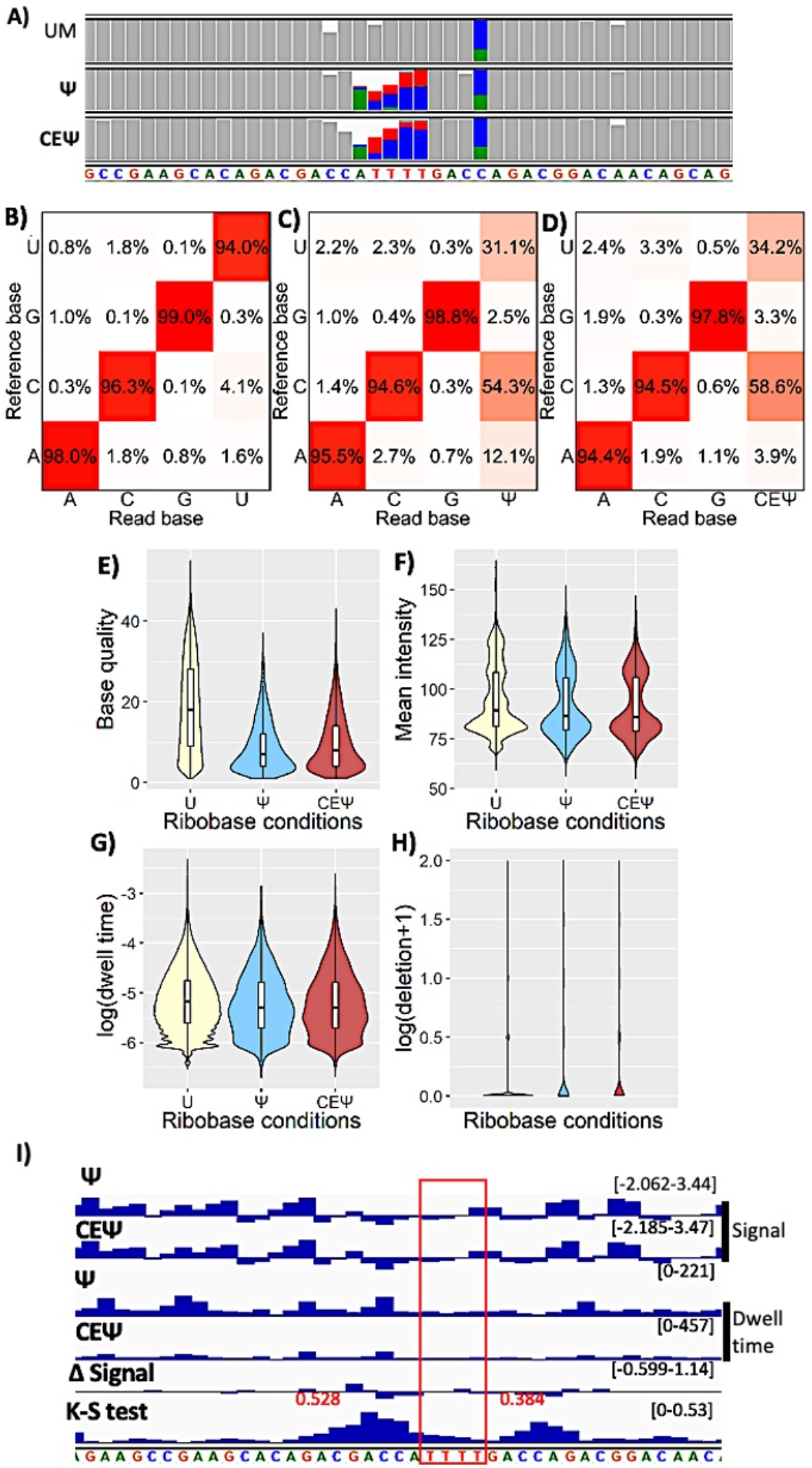
Altered nanopore parameters by Ψ and CEΨ modified nucleobases. **A)** IGV snapshot of unmodified, pseudouridine (Ψ) and cyanoethylpseudouridine (**CEΨ**) transcripts showing mismatch. Mismatch frequency > 0.2% are represented in colors. Green(adenosine), orange (guanosine), blue (cytosine) and red (thymine). **B, C & D**) Substitution matrix of unmodified, **Ψ** and **CEΨ** transcripts native reads, respectively. The x-axis represents the base identity of nanopore reads. The y-axis represents base identity of reference transcript **E-H**) Violin plot showing kernel density estimate & inner boxplot showing interquartile range and median of unmodified, **Ψ** and **CE Ψ** nucleobase. **E**) Base quality, **F**) Mean intensity, **G**) Dwell time and **H**) Deletion parameters. **I**)IGV snapshot showing signal comparison between **Ψ** (control) and **CEΨ** (samples) using TOMBO. The K-S test was used to compare signal difference (Δ signal) between **Ψ** and **CEΨ** conditions. The K-S test value > 0.2 is considered to harbor statistically significant signal difference.

We then tested our chemical probe induced mismatch error-based dRNA-Seq method on complex *in vivo* transcriptome using ribominus mouse brain RNA. We generated the following sample conditions, 1) untreated dataset referred to as **CE (-)** and 2) acrylonitrile-treated referred to as **CE (++)** (**Figure 4A**). Before dRNA-Seq, the acrylonitrile reactivity on mouse transcriptome was confirmed using cyanoethylation coupled with Sanger sequencing, a gold standard method for **A**-**I** validation. As expected, the cyanoethylation mediated depletion of **I** peak was observed in the bonafide **A**-**I** editing sites on Gria2, Kcan1 gene (**Figure S9**)^20^. All the samples were sequenced using independent flow cells for mapped read across conditions (**Table S1)**. The dRNA-seq resulting FAST5 files were basecalled to FASTQ files using GUPPY, which were further mapped using Minimap2 against the mouse genome/transcriptome references (refer methods). Good replicability (r = 0.93-0.95) was observed in terms of gene count across three conditions indicating that the acrylonitrile treatment did not alter the dRNA-Seq coverage **(Figure 4B, C**). Owing to the high reactivity of acrylonitrile towards **I** than Ψ, we primarily focused only on **I** in the transcriptome analysis. We retrieved chromosome coordinates of **A**-**I** sites through RADAR^20^, REDI portal database^21^ and published literatures^22^. From the above **A**-**I** sites, we further removed sites overlapping with the latest version of the SNP database (dSNP 142). These high confidence pre-annotated sites overlapped with our mouse dRNAseq, with a read count cut-off of > 5 yields thirteen high-confident **A**-I sites with mismatch errors **(Table S2)**. To obviate the possible error during library preparation, we assessed the read count correlation between our dataset and a previously published mouse brain dRNA-Seq dataset^23^. A high correlation (r=0.84-0.87) between two independent datasets **(Figure S10)** suggests that the lesser number of **I** sites recovered from our analysis could be due to the well-known coverage limitations associated with the long-read sequencing.

**Figure 4.**
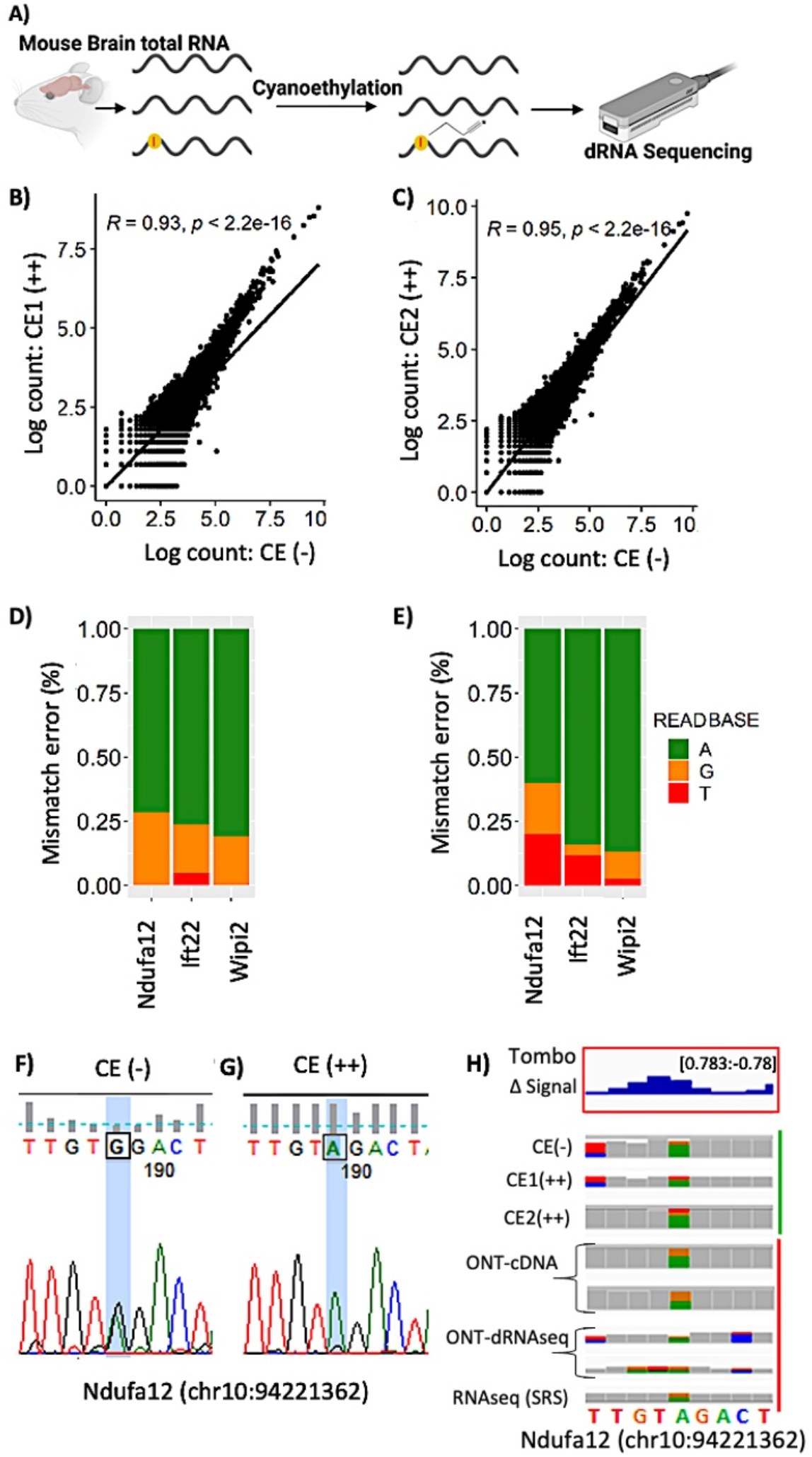
Validation of chemical probe-based nanopore sequencing method for the detection of RNA modifications. **A)** Overview showing different conditions of mouse brain samples subjected to sequencing. **B&C**) Agreement between different conditions of mouse brain dRNAseq. **B)** CE (-) versus CE1(++) and **C)** CE (-) versus CE2(++). **D&E**) Barplot showing mismatch frequency on pre-annotated inosine modification sites **D)** CE (-) and **E)** merged reads from CE1(++) and CE2(++). **F&G)** sanger validation of inosine sites through cyanoethylation based inosine chemical erasing. **F)** CE (-) conditions showing A/G dual peak, where G peak resulting from A-I modification **G)** CE (++) condition showing depletion of G peak, indicating true inosine modification. Traces for A, G, T and C are coloured in green, black, red and blue, respectively. **H)** IGV snapshot showing signal difference (TOMBO) and mismatch error in Ndufa12(chr10:94221362) gene. Green and red side annotations of IGV tracks represent mouse brain dRNAseq produced in this study and a public dataset, respectively.^25^

Most **A**-**I** sites captured in our mouse dRNA-Seq are less than the default threshold of 50 reads recommended by TOMBO for statistical analysis. Based on the read counts criteria ranging from highest to lowest, we selected four out of thirteen sites and performed Sanger sequencing validation to confirm the **I** modification sites inferred from dRNA-Seq mismatch errors (**Figure 4D**). Consistent with our notion, the dRNA-Seq mismatch error analysis coupled with sanger validation indicated that three sites (Ndufa12, Wipi2 and Ift22) harbored **A-I** modification (**Figure 4F-H**). However, the dRNA-Seq signature mismatch error inferred as the **I** in the Tox4 site was revealed to harbor SNV (**Figure S11C**). Three **A-I** sites validated by Sanger sequencing displayed a differential mismatch error pattern between CE (-) and CE (++) conditions. Besides, **A-I** modification abundancy among three sites from sanger validation falls in the following order Ndufa12 > Wipi2 > Ift22 **(Figure S11)**. It is important to note here that the hyper **A**-**I** editing of Wipi2 transcripts was reported in the prefrontal cortex of chronic social defeat stress susceptible mice^24^. NduFa12 ^25^and Itf22 are also associated with neurodegenerative disorders. Similar to **A**-**I** modifications, we overlapped the CeU-Seq^26^ captured **Ψ** sites with our mouse transcriptome reads and observed twenty-seven sites harboring signature mismatch errors attributable to **Ψ** modification (**Figure S12, Table S3**). Similar to the pattern observed in the synthetic **Ψ** modified RNA, the mismatch error profile is between **C** and **U**. Therefore, our results indicate that chemically induced mismatch errors can facilitate the selective assessment of RNA modifications.

In this proof-of-concept study, we have demonstrated the ability of the selectively reactive chemical probe like acrylonitrile to complement dRNA-Seq for profiling epitranscriptome with single-nucleotide resolution. The three modifications (**I, Ψ** and **m**^**6**^**A)** tested in this study were revealed by dRNA-Seq through their signature mismatch errors. Acrylonitrile treatment further enhanced the detection of **I-** and **Ψ** and distinguish it from other modifications with differential mismatch errors and signal difference. To prove this methodology’s applicability, we validated their detection ability at the transcriptome-wide level in mammalian mouse brain RNA. In the *in vivo* analysis, the acrylonitrile adducts revealed **I** modification even without the null data set. Furthermore, **A-I** modifications captured at the selective sites using dRNA-Seq were validated by sanger sequencing. While the low coverage is a major limitation in the dRNA-Seq platforms, the rapid developments in dRNA-Seq sequencing depth, accuracy and new modification-specific analysis pipelines could improve their accuracy. Our work opens up a new avenue of identifying other RNA modifications using selectively chemical probes^27,28,29^and chart the uncharted epitranscriptome landscape. Hypo or Hyper editing landscape of RNA modifications are reported in different disease conditions. Specifically, **A**-**I** differential editing has been reported in various psychiatric disorders such as autism, schizophrenia, and epilepsy^30^. However, such epitranscriptome changes were not explored for diagnostic purposes due to the lack of “Quick to Adapt” profiling methods. ONT platform-based epitranscriptome profiling can be easily developed as a clinically deployable diagnostic tool. However, generating a comparative dataset is an issue as gene knockout cannot be performed in humans. Chemical probe-based strategy could reliably identify RNA modification without the need for a null-modification dataset. Also, multiple chemical probes against various RNA modifications could facilitate simultaneous identification of different modifications. A recent study demonstrated that isoform resolved RNA structure in the dRNA-Seq platform using chemical probes^31^. Future studies combining probes for RNA modification and structure could accelerate our understanding of how epitranscriptome and RNA structure co-regulate gene expression at the transcriptome-wide level.

## AUTHOR CONTRIBUTIONS

S.R and V.J.S. contributed equally to this work. S.R. conceived the idea. G.N.P, S.R and V.J.S. designed the work. S.R and V.J.S. performed research, S.R. analyzed the data. L.C., T.H., Y.Z. and H.S. gave critical comments to improve the work. S.R, V.J.S. and G.N.P. wrote the paper. The authors declare no conflict of interest.

## ACKNOWLEDGEMENTS

This work was supported by Japan Society for the Promotion of Science (JSPS) KAKENHI (JP19H03349 to G.N.P, Grant No. JP16H06356 to

H.S and AMED under Grant No. JP18am0301005 (Basic Science and Platform Technology Program for Innovative Biological Medicine), JP20am0101101 (Platform Project for Supporting Drug Discovery and Life Science Research (BINDS)). We thank MEXT fellowship for V.J.S. We also thank JSPS #S19127; NIH/NEI EY031439-01; NJCSCR grant #151RG006 and Rutgers - TechAdvance Fund.

## DATA AVAILABILITY

All FASTQ files generated in this work are publicly available at European Nucleotide Archive (ENA).

## Supporting Information

## MATERIALS AND METHODS

### IVT template design and synthesis

Double-stranded DNA templates with T7 promoter for synthetic RNA were commercially purchased from IDT as blocks, which also has poly-A tail for nanopore adapter binding. IVT reactions were performed using MEGA script™ T7 Transcription Kit (AM1334), for overnight at 37°C followed by DNase treatment and purified using Quick Spin Columns for radiolabelled RNA purification (Roche, 11274015001). For **Ψ** and **I** modified RNAs synthesis, Pseudouridine-5’-Triphosphate (N-1019, Trilink) and Inosine-5’-Triphosphate (N-1020, Trilink) were used in place of uridine and guanosine, respectively.

### Inosine IVT reactions

Inosine IVT RNA was synthesized as two variants, 100% - **I** and 50%-I modified. 100% - **I** modification containing transcripts were synthesized using four ribonucleotides, G-cap analog (m7G(5’)ppp(5’)G, Invitrogen) , ITP, ATP, CTP and UTP. 50% I modified were synthesized using five nucleotides 50% GTP, 50% ITP, ATP, CTP and UTP.

### Acrylonitrile cyanoethylation reaction

CE solution is prepared with 5.0 ml of 100% ethanol, 1.53 ml of 7.19 M triethylamine (34804-85, Nacalai), and the pH was adjusted to 8.6 using acetic acid. The cyanoethylation of modified RNAs used 500ng of RNA, 30 µl of CE solution and 1.6 M of acrylonitrile and incubated at 70°C for 30 mins. Then, the reaction was immediately quenched by adding 160 ul of nuclease-free water on ice. Cyanoethylated RNA was precipitated using 0.1 volume of 3M sodium acetate and three volumes of 100% ethanol.

### HPLC and Mass spectrometry validation of cyanoethylation reaction

For nucleoside analysis, the IVT template RNA (0.01– 0.05 A260 units) was digested into nucleosides using 10 µg/ml nuclease P1 (New England Biolabs, M0660S), and 0.5 U/ml Bacterial Alkaline phosphatase (Takara 2120A) at 37°C for 1 h in 10 µl of a reaction mixture containing 20 mM HEPES–KOH pH 7.5(Nacalai 15639-84). The nucleosides were separated using an HPLC equipped with Chemcobond 5-ODS-H column (4.6 × 150 mm). Analysis was performed with a mobile phase with solvent A containing 0.1% TFA (Trifluoracetic acid) in water in gradient combination with solvent B of acetonitrile. The linear gradient started with 0% solvent B at 0 min to 10% B for 30 min, at a flow rate of 1 mL min^−1^, monitored at 254 nm. The exact mass was confirmed using a Q Exactive-Orbitrap mass spectrometer (Thermo Fisher, Germany).

### Direct RNA library preparation and sequencing

500ng of RNA was used for direct RNA seq library (SQK-RNA002) preparation following the ONT protocol version - DRS_9080_v2_revK_14Aug2019. Briefly, 500 ng of unmodified, modified and CE treated IVT RNA were ligated to ONT RT Adapter using concentrated T4 DNA ligase (NEB-M0202T), and was reverse transcribed using SuperScript III Reverse Transcriptase (Thermo Fisher Scientific-18080044). The products were purified using 1.8X Agencourt RNAClean XP beads (Fisher Scientific-NC0068576), washing with freshly prepared 70% ethanol. RNA Adapter (RMX) was ligated onto the RNA: DNA hybrid and the mix were purified using 1X Agencourt RNAClean XP beads, washing with wash buffer twice. The sample was then eluted in elution buffer and mixed with RNA running buffer (RRB) prior to loading onto a primed R9.4.1 flow cell and ran on a MinION sequencer with MinKNOW acquisition software version v1.14.1.

### Mouse brain RNA isolation and dRNAseq

Three months old ICR mice were purchased from (Shimizu, Japan). Forebrain was dissected and homogenized using Dounce homogenizer in 2ml of sepasol-RNA I super G (Nacalai Tesque). To the 1 ml of homogenized sample 200ul of chloroform was add. After centrifugation at 13,000g for 15 mins at 4 °C, the aqueous layer was transferred and combined with an equal volume of 70% ethanol. RNeasy column with DNase I treatment used for RNA purification according to the manufacturer’s instructions (RNase Plus Mini Kit, Qiagen). Ribominus RNA was prepared using RiboMinus™ Eukaryote Kit v2 (Invitrogen). Minimum 500ng of Ribominus RNA is subjected to CE treatment. Integrity and concentration of RNA was checked using Bioanalyzer 2100 RNA Nano kit (Agilent), following standard protocols. After CE treatment we mostly recovered > 400ng of mouse brain RNA which was subjected to dRNAseq library preparation as mentioned above.

### Base calling and mapping

FAST5 files were basecalled using MinKnow-GUPPY (V 3.4.5) with accurate base calling enabled. Mapping to reference sequence were done using minimpa2 (version-2.17-r941) with the setting -ax map-ont -L. Mapped reads were further filtered to remove unmapped, secondary, and supplementary and reads mapped on the reverse strand. Reads with low alignment were also removed with cut off MAPQ < 10, using options -bh -F 2324 -q 10. These reads were further sorted and indexed. This mapping workflow is adapted from https://nanocompore.rna.rocks/. Reference file used to map IVT reads are given in the supplementary file. Mouse brain reads were mapped to gencode. vM24 transcript reference (release M24) and reference genome (release Grcm38.p6). Read counts of mouse dRNA-seq datasets were extracted using salmon (V1.4.0). Salmon quantification requires transcript reference mapped bam files.

### Extraction of nanopore parameters

The nanopore parameters like base quality, mismatch, and deletion were extracted using scripts associated with epinano package (https://github.com/Huanle/EpiNano). The epinano scripts were used to extract the per-site feature of the above-described parameters.

- samtools view -h file.bam| java -jar sam2tsv.jar -r ref.fasta > file.bam.tsv
- python2 per_read_var.py file.bam.tsv > file.per_read.var.csv The dwell time and current intensity of k-mers were extracted using scripts associated with https://nanocompore.rna.rocks/ using Nanopolish (v 0.11.1) and NanopolishComp (v.0.6.11)
- nanopolish index -s {sequencing_summary.txt} -d {raw_fast5_dir} {basecalled_fastq}
- nanopolish eventalign --reads {basecalled_fastq} --bam {aligned_reads_bam} --genome {transcriptome_fasta} -- samples --print-read-names --scale-events --samples > {eventalign_reads_tsv}
- NanopolishComp Eventalign_collapse -i {eventalign_reads_tsv} -o {eventalign_collapsed_reads_tsv}

All the plots in this paper ate generated using R (4.0.2)

### Tombo analysis (level sample compare)

The input for TOMBO (v1.5.1) analysis(https://nanoporetech.github.io/tombo/) is single FAST5 format. Current ONT runs result in muti-Fast5 format, first it was covert to single-FAST5 format using **multi_to_single_fast5** (v3.0.1). Next FAST5 files were annotated with respective FASTQ files using **tombo preprocess annotate_raw_with_fastqs**. After above step **tombo resquiggle** is employed to align mapped reference location and raw signal to the sequence alignment. Above steps were independently followed for both CE (-) and CE (++) conditions, then both the conditions were compared using **tomb detect_modifications level_sample_compare**. IGV genome browser compatible files are further extracted from the **tombo detect_modification** generated statistics file in using **tombo text_output browser_files**.

**Figure S1.**
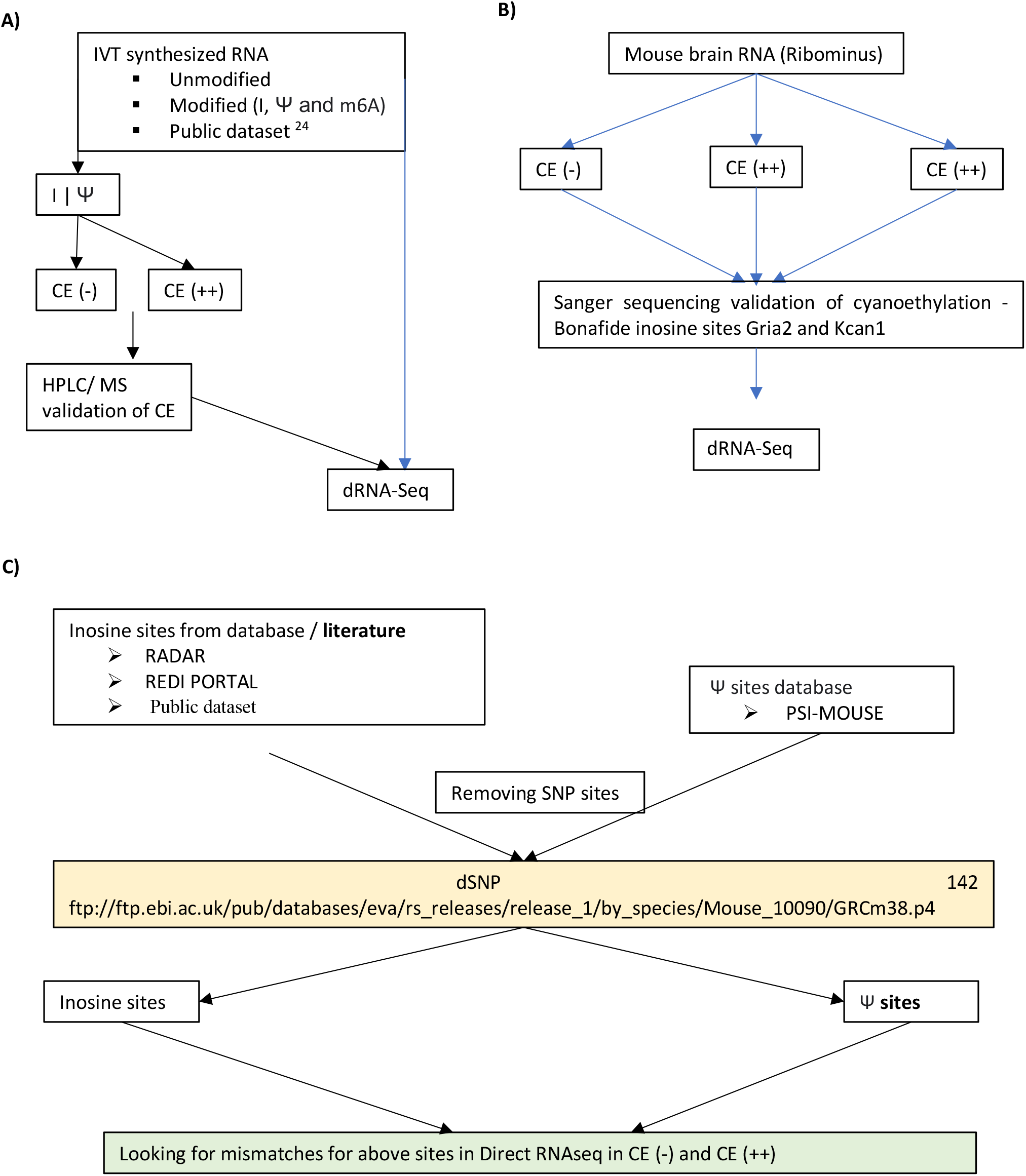
Schematics of experiments and analysis workflow. **A)** IVT RNA preparation **B)** mouse brain RNA preparation and, **C)** analysis pipeline employed to score pre-annotated **I**/ **Ψ** sites from mouse dRNAseq reads

**Figure S2.**
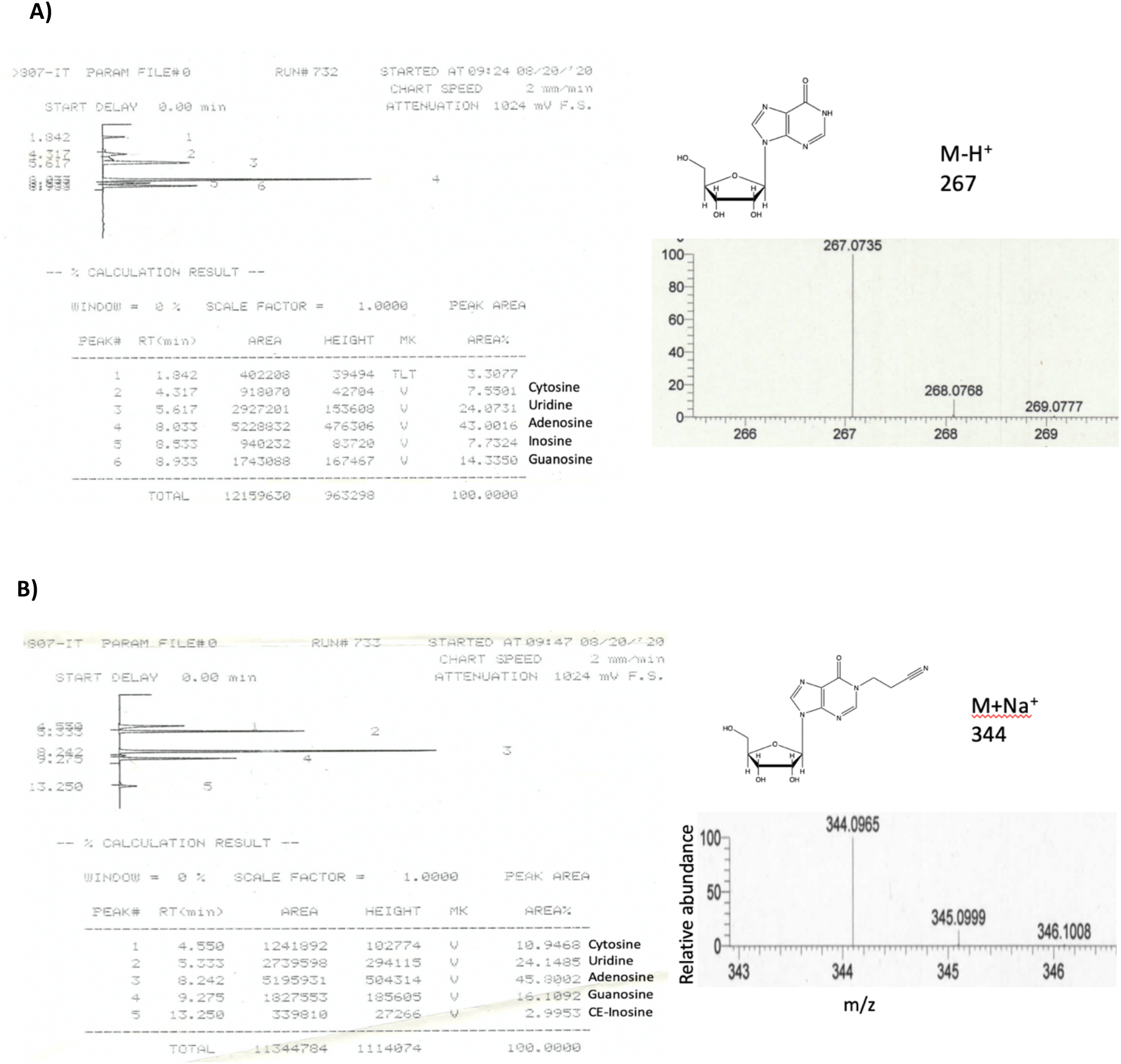
HPLC of digested IVT RNA and mass spectrometry corresponding to the molecular weight of **A)** Inosine and **B)** CEI

**Figure S3.**
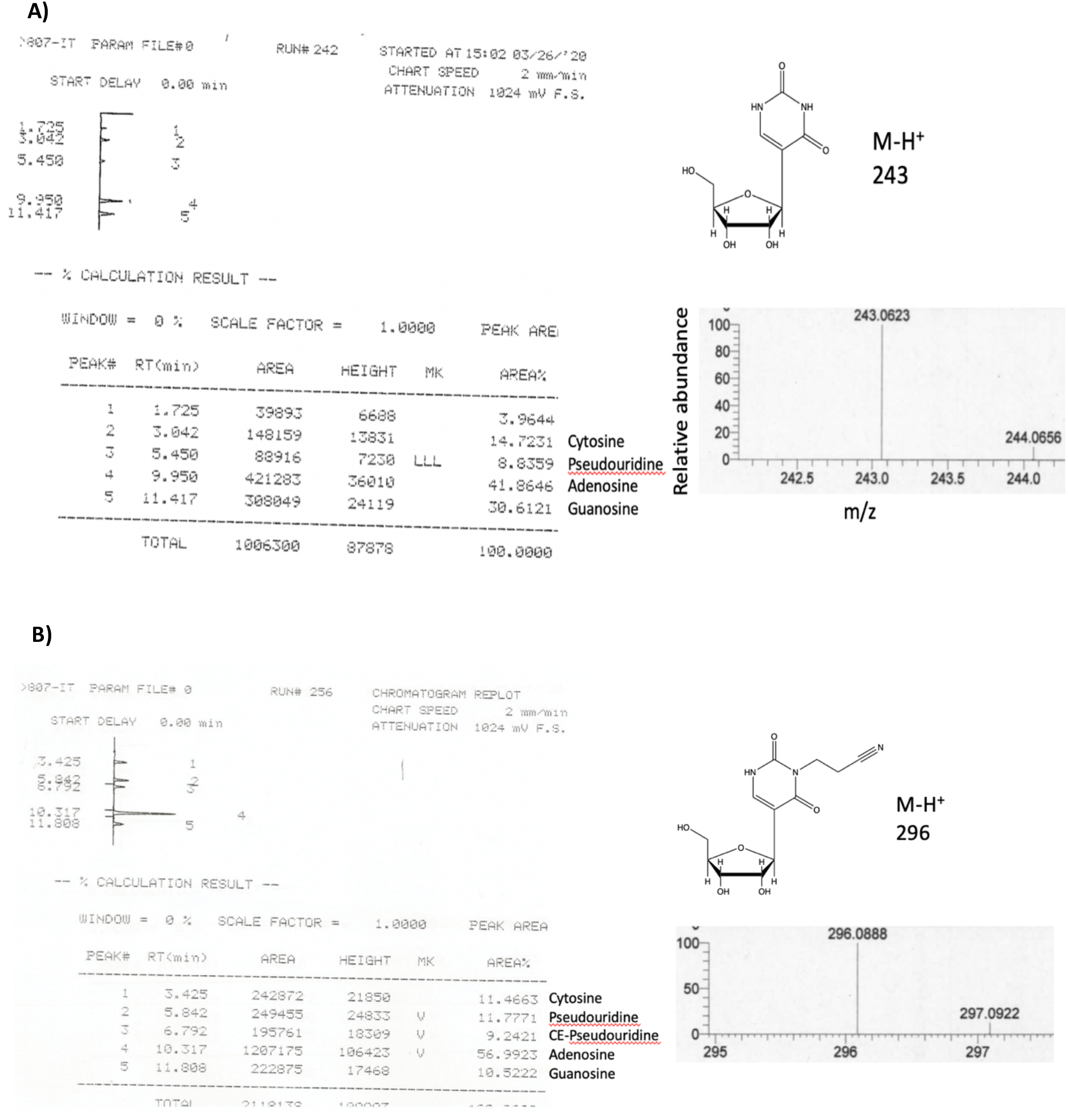
HPLC of digested IVT RNA and mass spectrometry corresponding to the molecular weight of **A) Ψ** and **B) CEΨ**.

**Figure S4.**
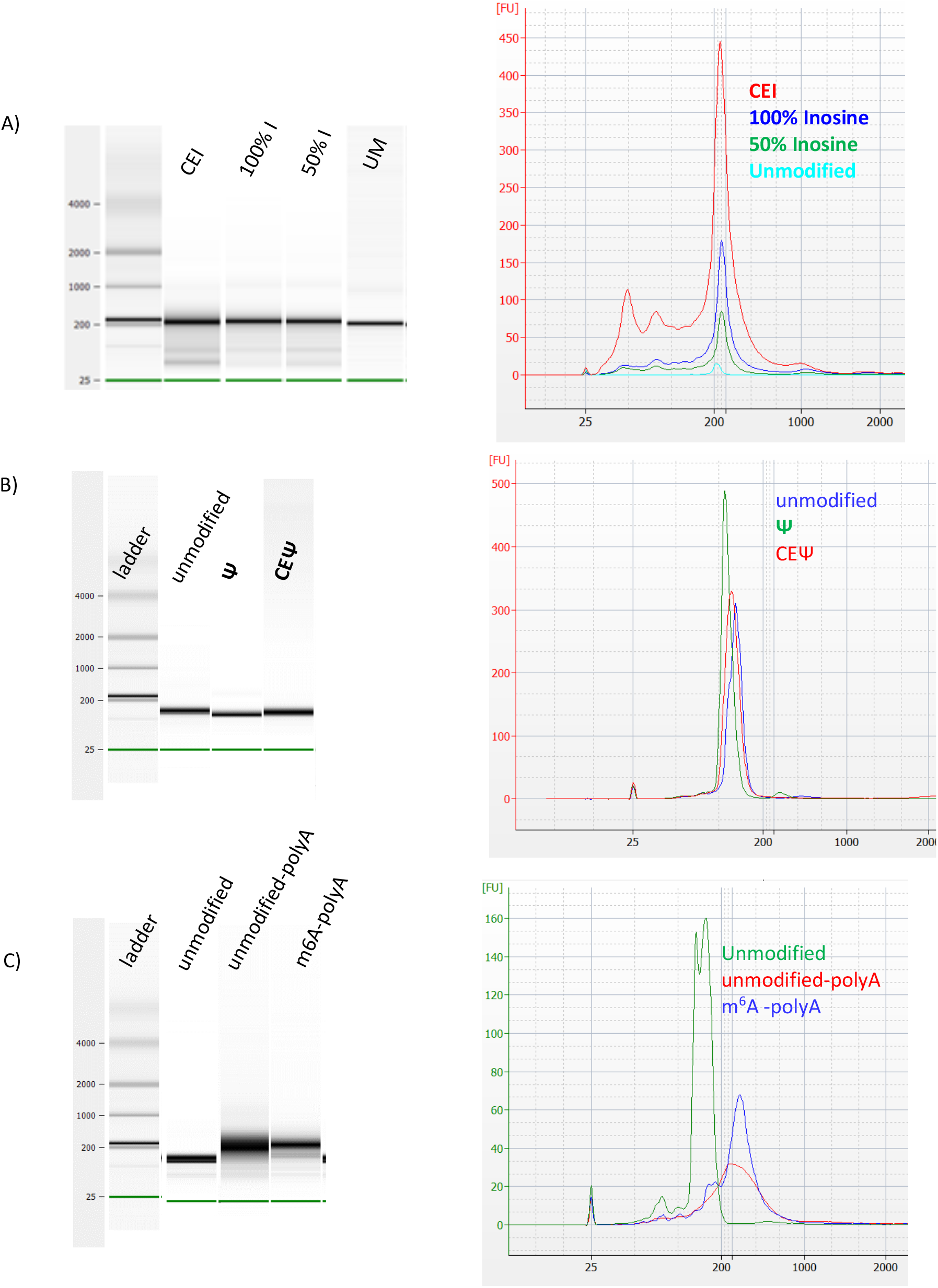
Bioanalyzer of IVT RNA integrity show gel picture and electropherogram **A)** Inosine, **B) Ψ**, & **C) m**^**6**^A

**Figures S5.**
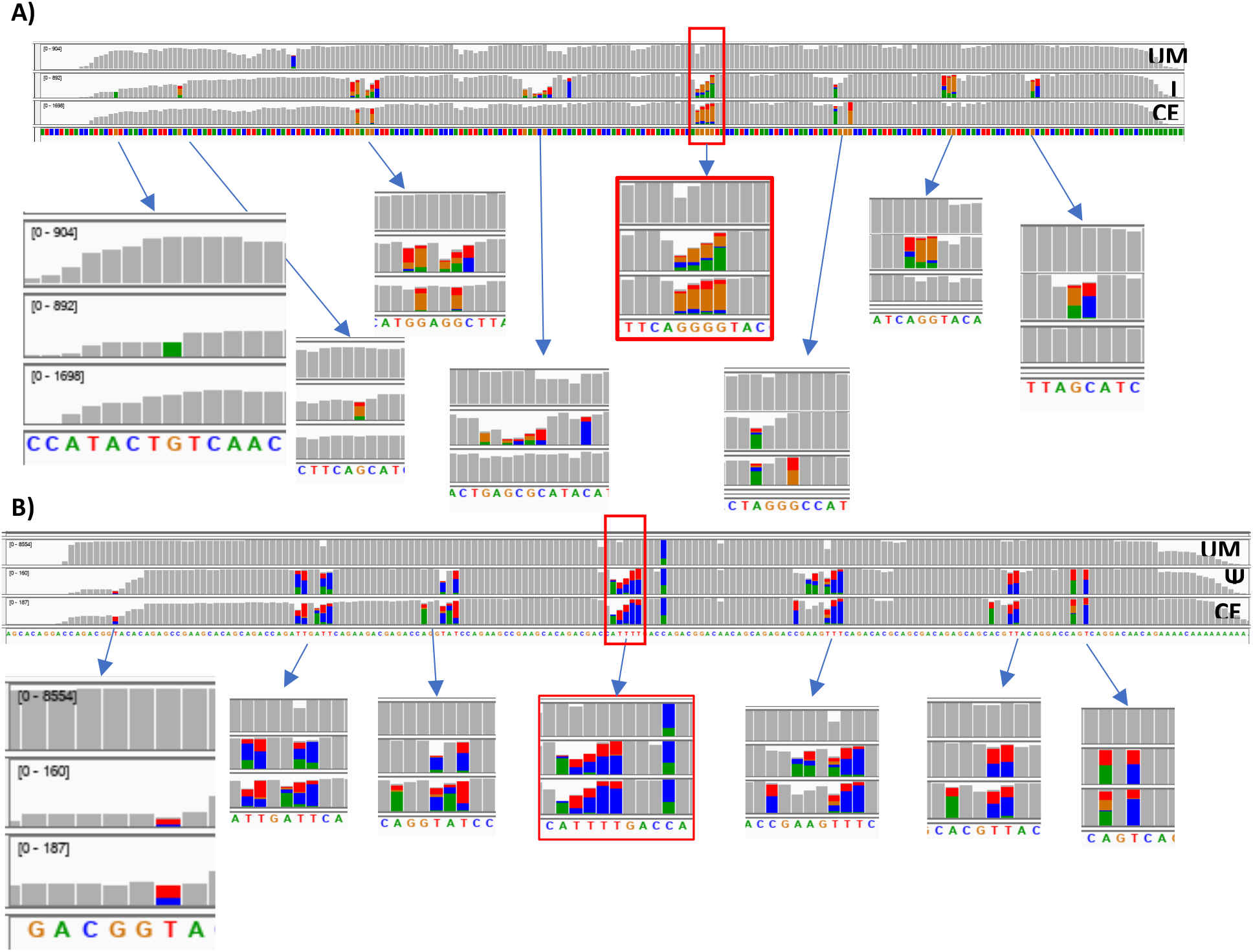
IGV snapshot showing mismatch errors in unmodified (UM), modified (**I, Ψ** and **m**^**6**^**A**) and cyanoethyl Inosine (**CEI**), cyanoethylpseudouridine (**CEΨ**) IVT RNA’s. **A)** Inosine, **B) Ψ**. Regions highlighted in main figures are indicated by red borders.

**Figure S6.**
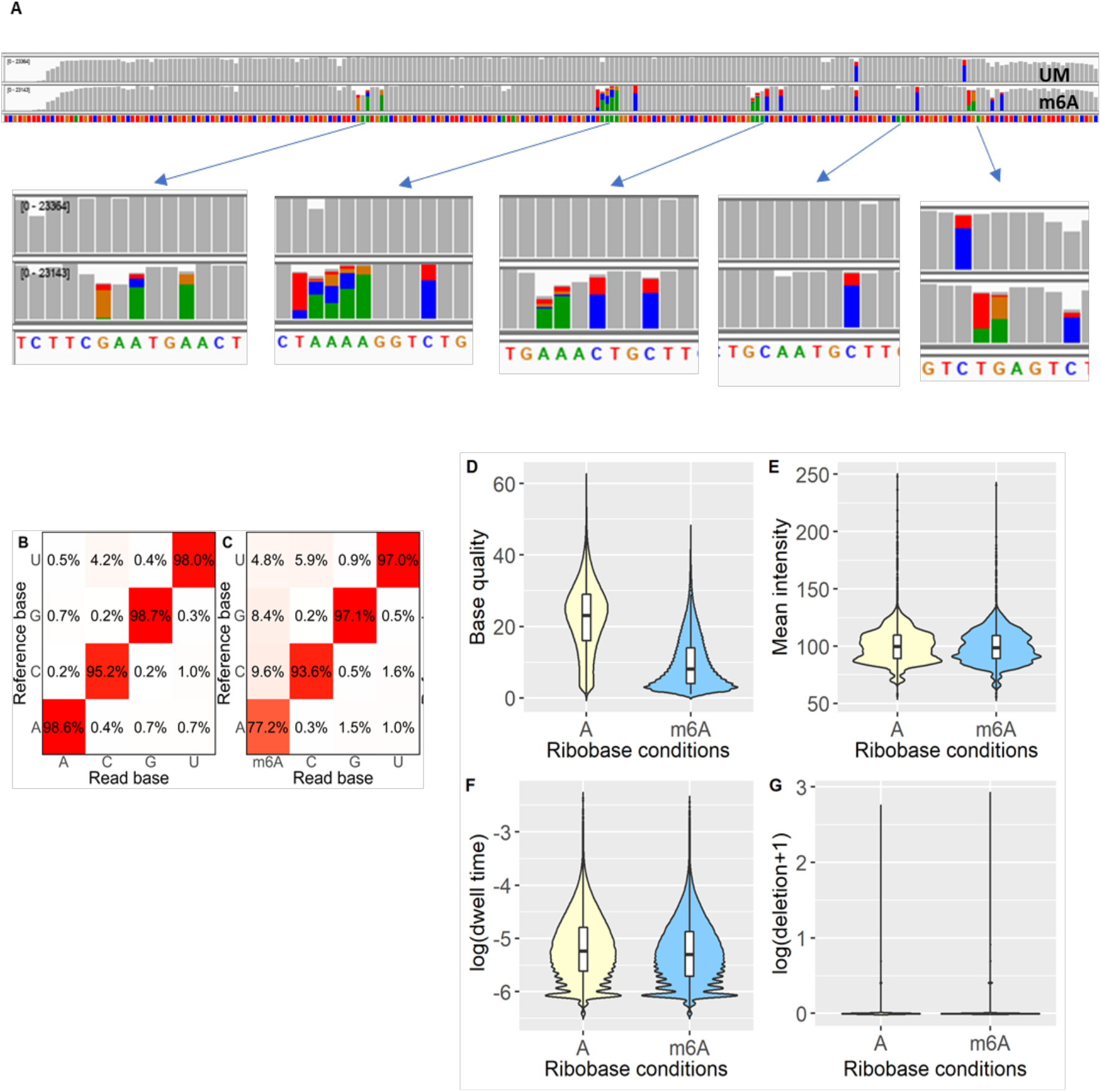
Altered nanopore parameters by **m**^**6**^**A** modified nucleobases. **A)** IGV snapshot of unmodified and **m**^**6**^**A** transcripts showing mismatch. Mismatch frequency > 0.2% are represented in colors. Green(adenosine), orange (guanosine), blue (cytosine) and red (thymine). **B&D)** Substitution matrix of unmodified and **m**^**6**^**A** transcripts native reads, respectively. The x-axis represents the base identity of nanopore reads. The y-axis represents base identity of reference transcript. **D-G)** Violin plot showing kernel density estimate & inner boxplot showing interquartile range and median of unmodified and **m**^**6**^**A** nucleobase. **D)** base quality, **E)** Mean intensity, F) dwell time and **G)** deletion parameters.

**Figure S7.**
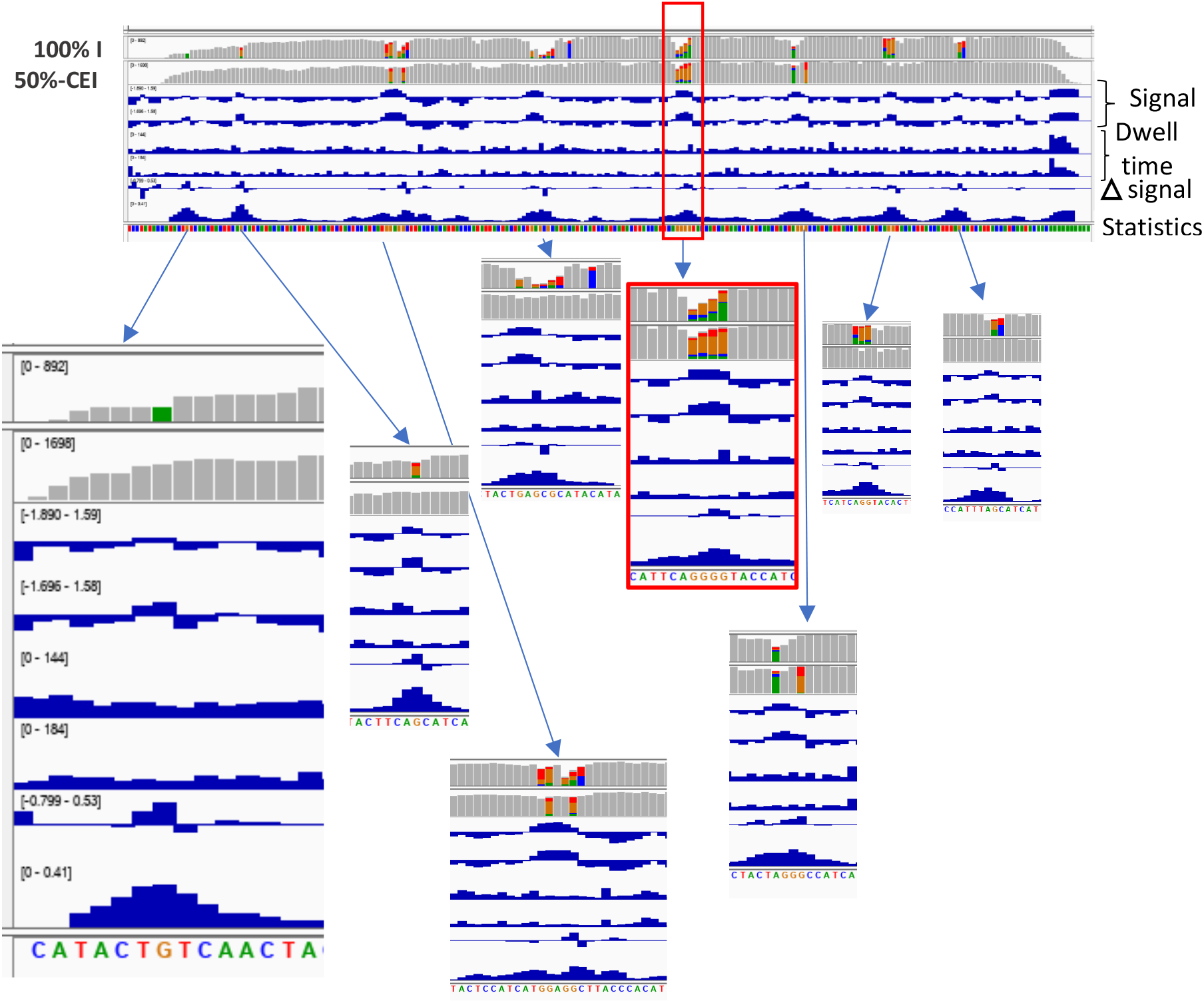
IGV snapshot showing signal comparison between **I** (control) and **CEI** (samples) using TOMBO. The K-S test was used to compare signal difference ((Δ signal) between **I** and **CEI** conditions. The K-S test value > 0.2 is considered to harbor statistically significant signal difference. Modified regions are further highlighted by zoom out panels, while the red border panel was also represented in the Figure 2I.

**Figure S8.**
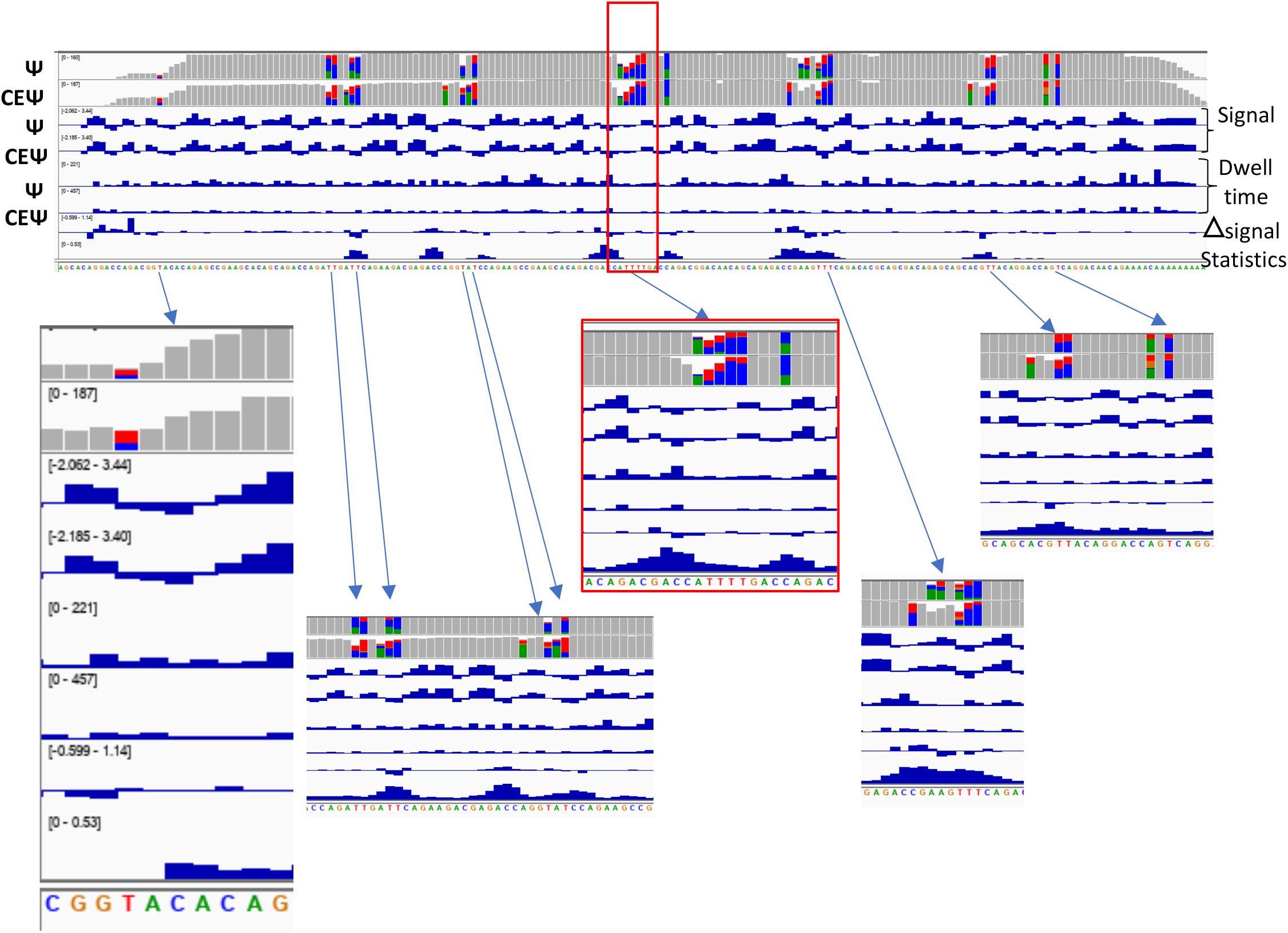
IGV snapshot showing signal comparison between **Ψ** (control) and **CEΨ** (samples) using TOMBO. The K-S test was used to compare signal difference ((Δ signal) between **Ψ** and **CEΨ** conditions. The K-S test value > 0.2 is considered to harbor statistically significant signal difference. Modified regions are further highlighted by zoom out panels, while the red border panel was also represented in the Figure 3I.

**Figure S9.**
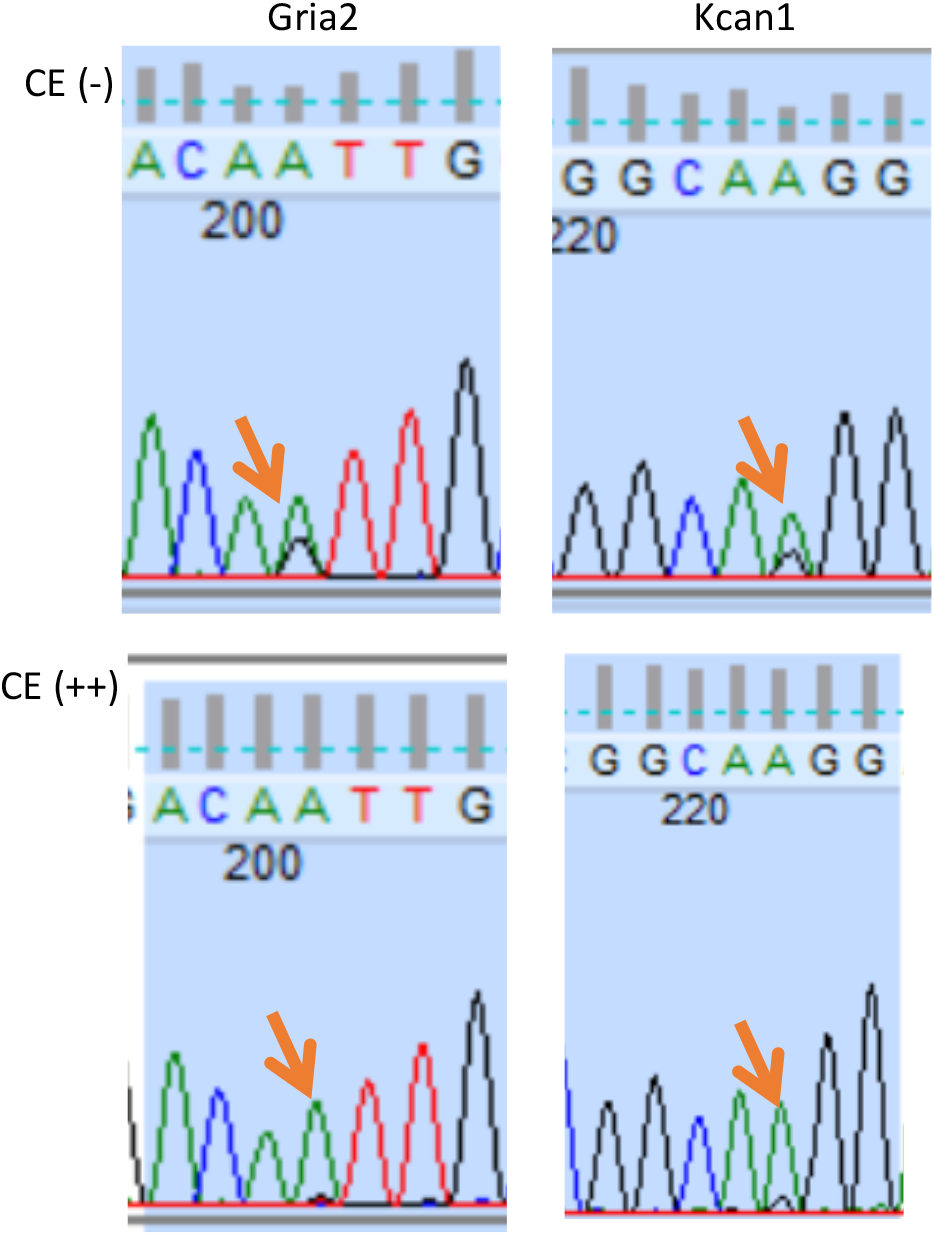
Sanger sequencing validates bonafide inosine sites in Gria2 and Kcan1 genes after cyanoethylation based inosine chemical erasing. CE(-) conditions showing A/G dual peak , where G peak resulting from **A**-**I** modification ,CE(++) condition showing depletion of G peak , indicating true inosine modification. Traces for A, G, T and C are coloured in green, black, red and blue, respectively.

**Figure S10.**
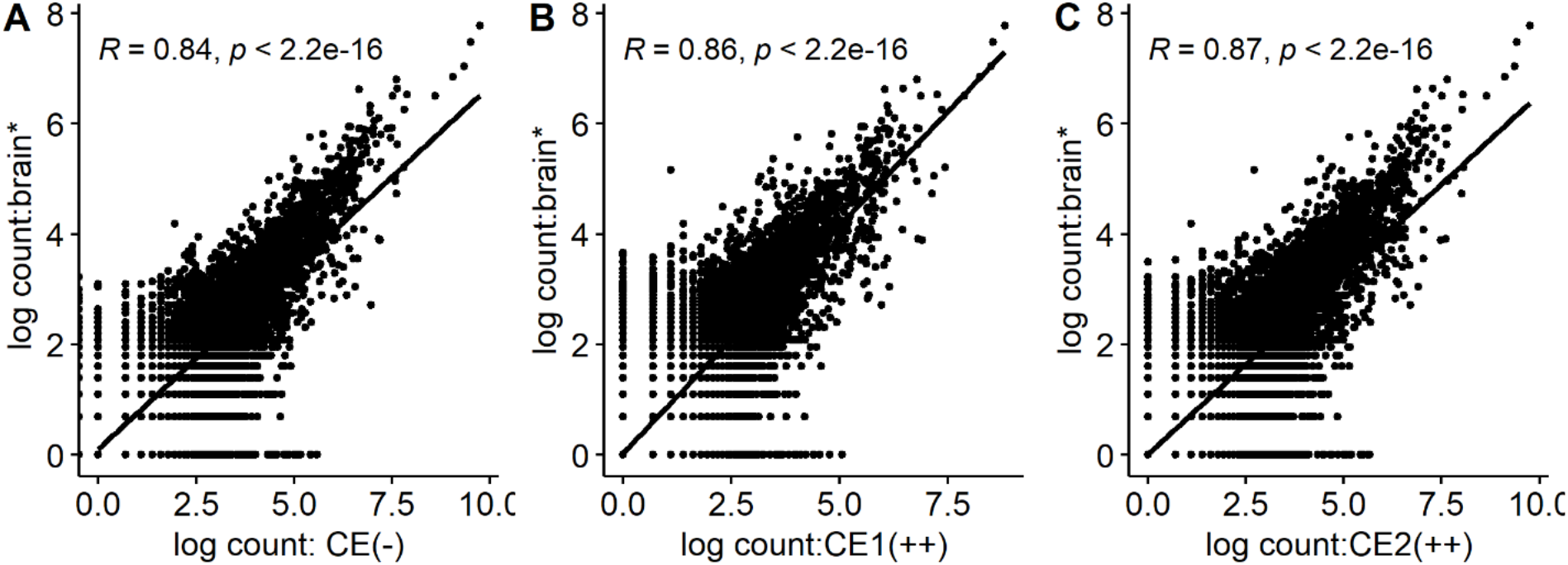
Correlation between read counts, over different conditions of our mouse brain dRNA-Seq versus public dataset ^25^ (brain*) dataset. A) CE (-) versus brain*, B) CE1(++) versus brain*, and C) CE2(++) versus brain*.

**Figure S11.**
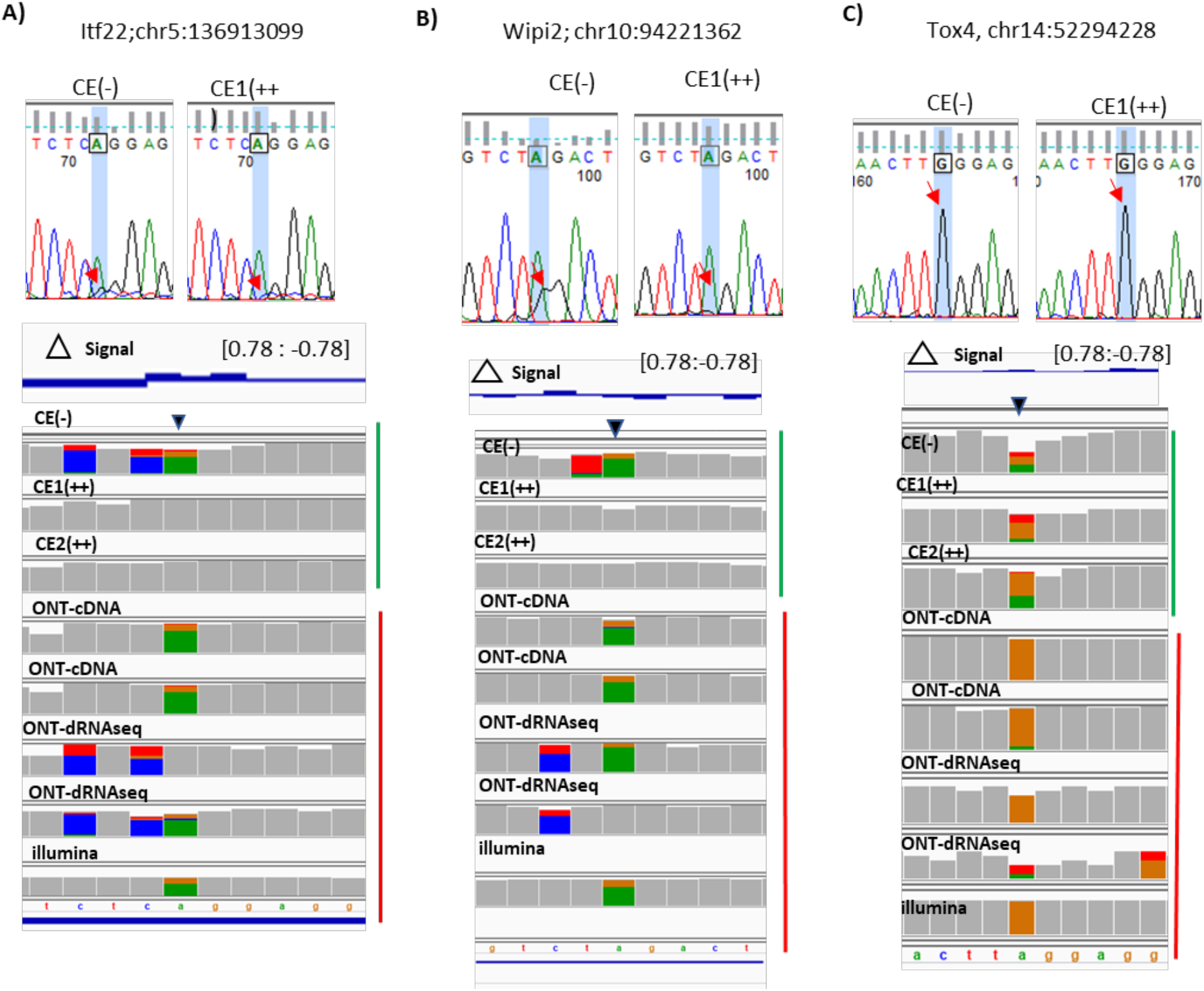
Sanger sequencing validates inosine sites through cyanoethylation based inosine chemical erasing. With CE (-), untreated conditions and CE (++) treated condition. A) Itf22, B) Wipi2 and C) Tox4.Traces for **A, G, T** and **C** are coloured in green, black, red and blue, respectively. Green and red side annotations of IGV tracks represent mouse brain dRNAseq in this study and from a public dataset, respectively.^25^

**Figure S12.**
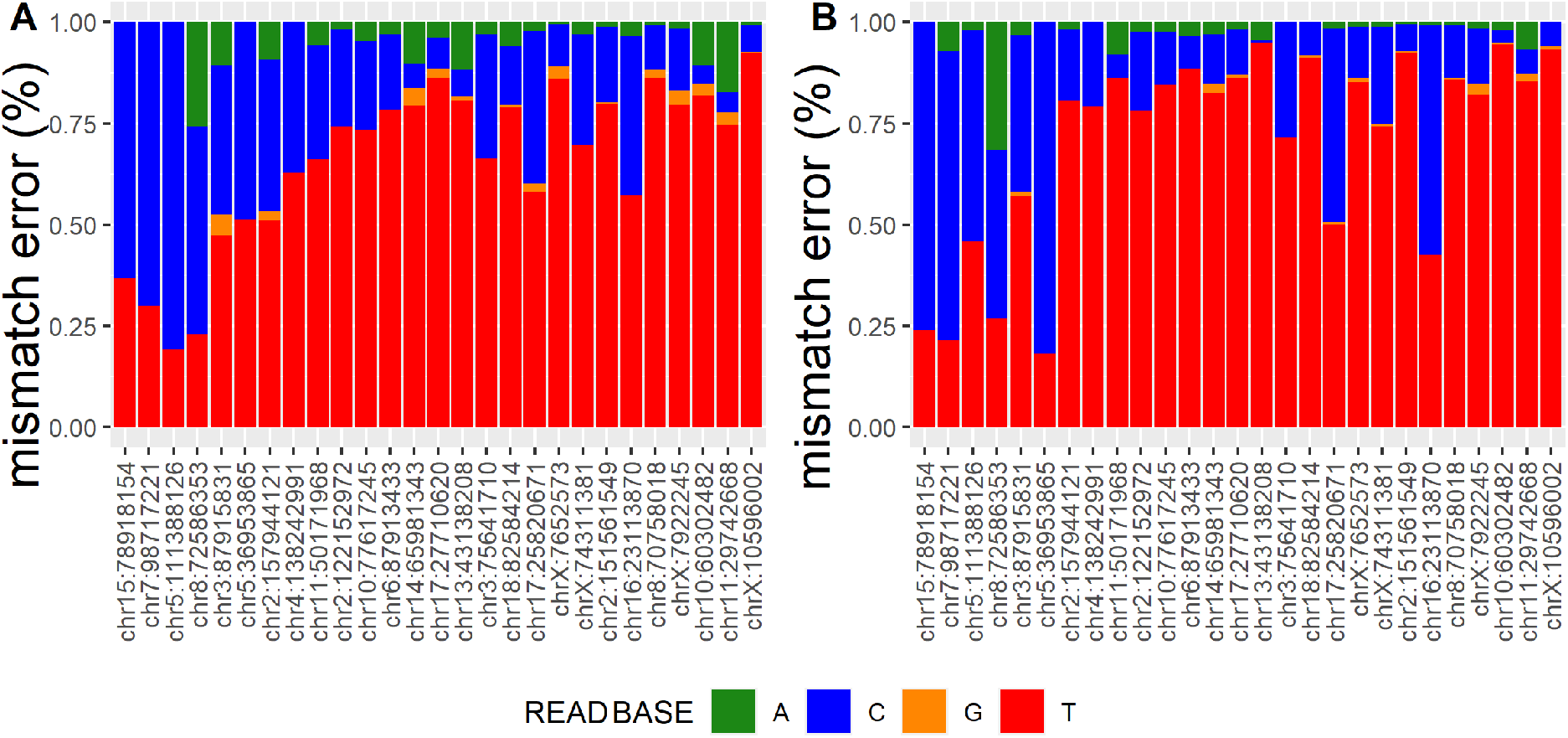
Barplot showing mismatch frequency on pre-annotated pseudouridine modification sites **A)** CE (-) and **B)** acrylonitrile treated CE (++).

## Supplementary File

### >Inosine (I)

AAGC<u>TAATACGACTCACTATAGG</u>ATCCTATACCATACTGTCAACTACTTCAGCATCATACACTACTTACATCATCTACTCCATCAT GGAGGCTTACCCACATTACCCATATTACTACTACTGAGCGCATACATACATCCATCATACTTACCATTCAGGGGTACCATCATAA CTCATCAACTACTAGGGCCATCATTACCATTCATCAGGTACACTTACCATTTAGCATCATTACCATCAATACAACAAAAAAAAAA

### >Pseudouridine (Ψ)

AAGC<u>TAATACGACTCACTATAGG</u>AGCACAGGACCAGACGGTACACAGAGCCGAAGCACAGCAGACCAGATTGATTCAGAAGA CGAGACCAGGTATCCAGAAGCCGAAGCACAGACGACCATTTTGACCAGACGGACAACAGCAGAGACCGAAGTTTCAGACACG CAGCGACAGAGCAGCACGTTACAGGACCAGTCAGGACAACAGAAAACAAAAAAAAAA

### > N6-methyladenosine (m6A)

aagc<u>TAATACGACTCACTATAGGGTCG</u>TGTTTCTCTTGTATGTCTGTCGTCTTGTCTCGCTGTGCTGCTGCTTCTGTCGTGCTTGC TTCGTTGCTCTTCGAATGAACTGTGCTTGCTGCTTCTGTTCTGCATAGCTGCTTCGTTCGTGCCTAAAAGGTCTGTCTCGTGCTCT GCTTGCTTGTGAAACTGCTTCGTCTGCTCTCGTCTTTGCTGCAATGCTTGTGTCGTCTGAGTCTCTTCGTCTCGTTTCGTGCTGC

### IVT RNA sequence design

T7 promoter regions are underlined.

Grey and red highlights the template sequence and modification positions, respectively.

**Table S1.**
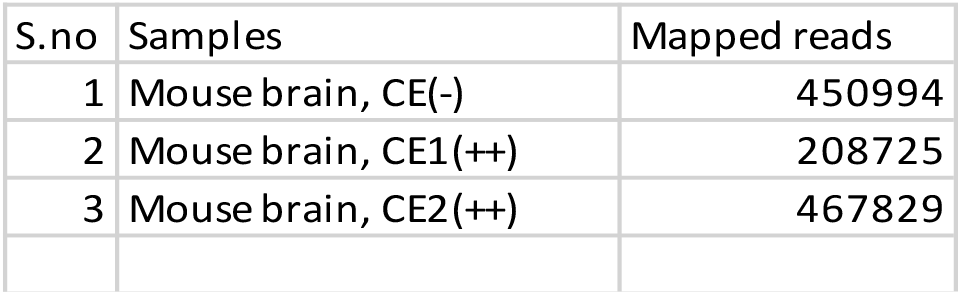
Mapped reads in mouse dRNAseq datasets. **Samtools-flagstat** was used to extract mapped reads from mouse dRNAseq dataset mapped to gencode transcript reference.

**Table S2.**
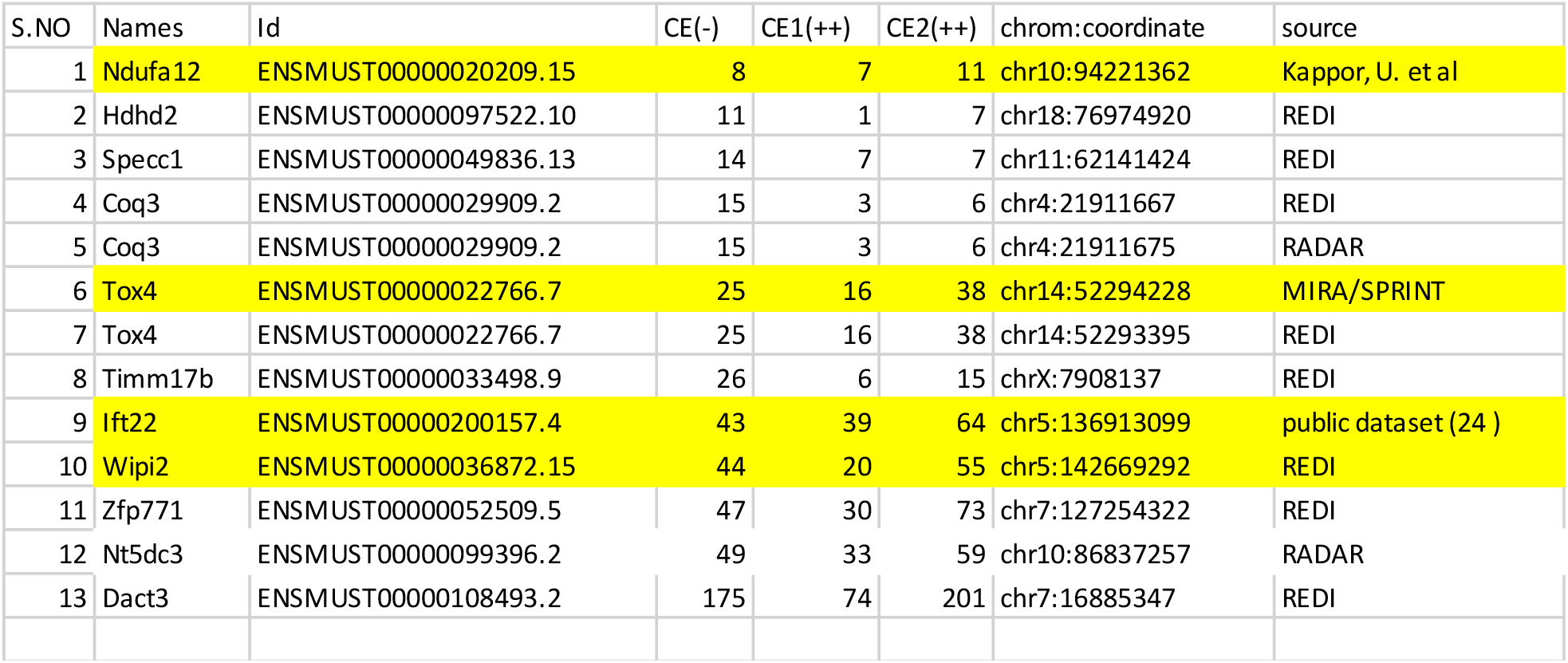
Read count of pre-annotated **A-I** modified sites present in mouse dRNAseq datasets. Yellow highlighted sites are validated using Sanger sequencing

**Table S3.**
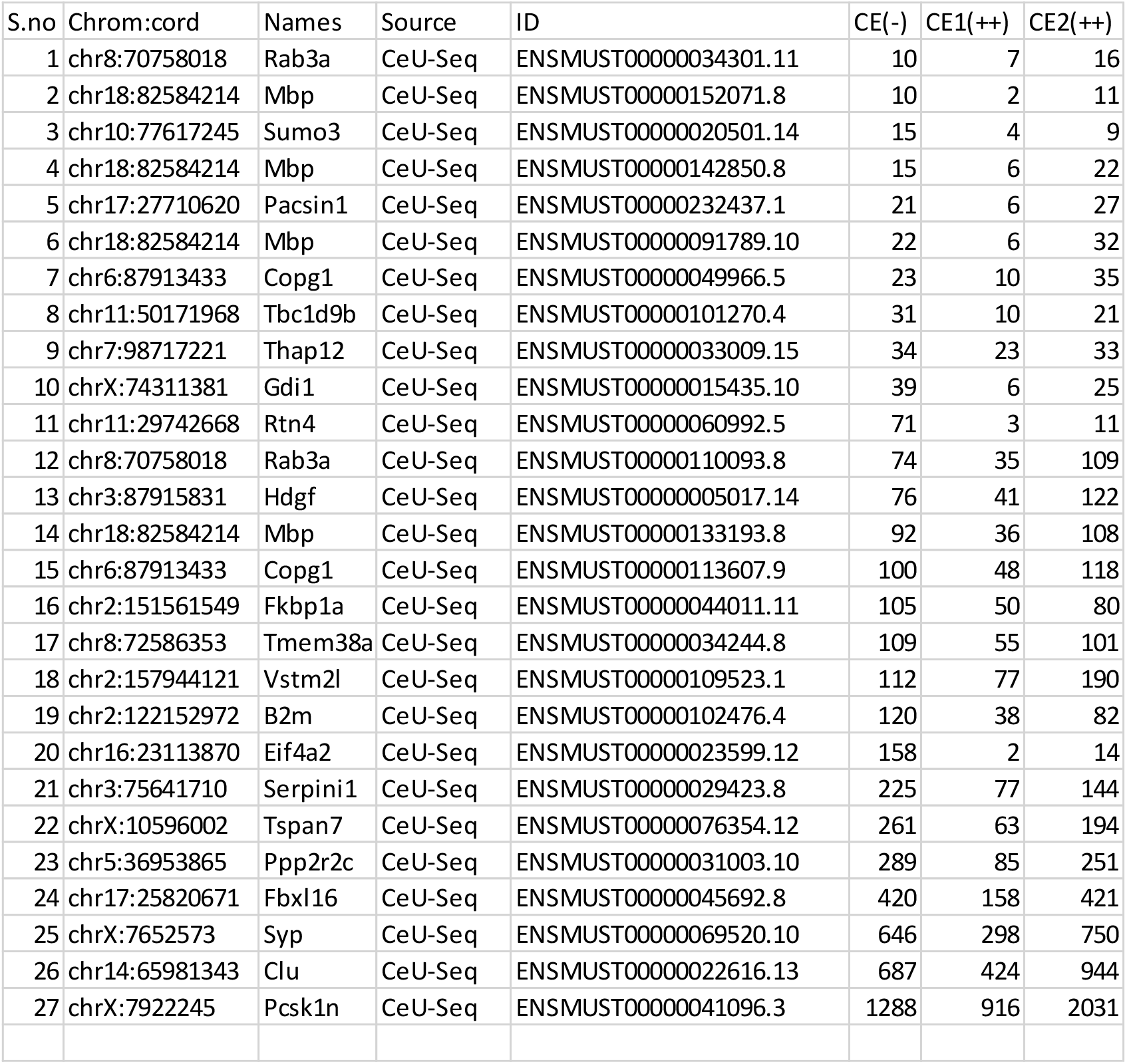
Read count of pre-annotated **Ψ** sites (CeU-seq) present in mouse dRNAseq datasets

| S.no | Chrom:cord | Names | Source | ID | CE(-) | CE1(++) | CE2(++) |
| --- | --- | --- | --- | --- | --- | --- | --- |
| 1 | chr8:70758018 | Rab3a | CeU-Seq | ENSMUST00000034301.11 | 10 | 7 | 16 |
| 2 | chr18:82584214 | Mbp | CeU-Seq | ENSMUST00000152071.8 | 10 | 2 | 11 |
| 3 | chr10:77617245 | Sumo3 | CeU-Seq | ENSMUST00000020501.14 | 15 | 4 | 9 |
| 4 | chr18:82584214 | Mbp | CeU-Seq | ENSMUST00000142850.8 | 15 | 6 | 22 |
| 5 | chr17:27710620 | Pacsin1 | CeU-Seq | ENSMUST00000232437.1 | 21 | 6 | 27 |
| 6 | chr18:82584214 | Mbp | CeU-Seq | ENSMUST00000091789.10 | 22 | 6 | 32 |
| 7 | chr6:87913433 | Copg1 | CeU-Seq | ENSMUST00000049966.5 | 23 | 10 | 35 |
| 8 | chr11:50171968 | Tbc1d9b | CeU-Seq | ENSMUST00000101270.4 | 31 | 10 | 21 |
| 9 | chr7:98717221 | Thap12 | CeU-Seq | ENSMUST00000033009.15 | 34 | 23 | 33 |
| 10 | chrX:74311381 | Gdi1 | CeU-Seq | ENSMUST00000015435.10 | 39 | 6 | 25 |
| 11 | chr11:29742668 | Rtn4 | CeU-Seq | ENSMUST00000060992.5 | 71 | 3 | 11 |
| 12 | chr8:70758018 | Rab3a | CeU-Seq | ENSMUST00000110093.8 | 74 | 35 | 109 |
| 13 | chr3:87915831 | Hdgf | CeU-Seq | ENSMUST00000005017.14 | 76 | 41 | 122 |
| 14 | chr18:82584214 | Mbp | CeU-Seq | ENSMUST00000133193.8 | 92 | 36 | 108 |
| 15 | chr6:87913433 | Copg1 | CeU-Seq | ENSMUST00000113607.9 | 100 | 48 | 118 |
| 16 | chr2:151561549 | Fkbp1a | CeU-Seq | ENSMUST00000044011.11 | 105 | 50 | 80 |
| 17 | chr8:72586353 | Tmem38a | CeU-Seq | ENSMUST00000034244.8 | 109 | 55 | 101 |
| 18 | chr2:157944121 | Vstm2l | CeU-Seq | ENSMUST00000109523.1 | 112 | 77 | 190 |
| 19 | chr2:122152972 | B2m | CeU-Seq | ENSMUST00000102476.4 | 120 | 38 | 82 |
| 20 | chr16:23113870 | Eif4a2 | CeU-Seq | ENSMUST00000023599.12 | 158 | 2 | 14 |
| 21 | chr3:75641710 | Serpini1 | CeU-Seq | ENSMUST00000029423.8 | 225 | 77 | 144 |
| 22 | chrX:10596002 | Tspan7 | CeU-Seq | ENSMUST00000076354.12 | 261 | 63 | 194 |
| 23 | chr5:36953865 | Ppp2r2c | CeU-Seq | ENSMUST00000031003.10 | 289 | 85 | 251 |
| 24 | chr17:25820671 | Fbxl16 | CeU-Seq | ENSMUST00000045692.8 | 420 | 158 | 421 |
| 25 | chrX:7652573 | Syp | CeU-Seq | ENSMUST00000069520.10 | 646 | 298 | 750 |
| 26 | chr14:65981343 | Clu | CeU-Seq | ENSMUST00000022616.13 | 687 | 424 | 944 |
| 27 | chrX:7922245 | Pcsk1n | CeU-Seq | ENSMUST00000041096.3 | 1288 | 916 | 2031 |

## Notes

### Competing Interest Statement

The authors have declared no competing interest.

### Summary of Updates

New Results updated

